# Preparation and characterization of Fe_3_O_4_-spermine-PCL-chitosan-PEG-FA nanoparticles for DNA delivery into AGS cells

**DOI:** 10.1101/2025.08.27.672642

**Authors:** Sahar Mohajeri, Negar Baezzat, Mehran Noruzpour, Shima Bourang, Hashem Yaghoubi

## Abstract

Gene therapy, a novel treatment approach for diseases such as cancer and genetic disorders, involves the transfer of genes to host cells or tissues. In recent decades, methods such as viruses, bacteria, and mechanical techniques have been utilized for gene transfer. However, Owing to the significant side effects of traditional methods for cancer treatment, especially chemotherapy, nanoparticles have emerged as a preferred alternative, enhancing targeted gene delivery and reducing complications. This study utilized copolymers such as spermine-chitosan-polycaprolactone (PCL) (PCS) and iron oxide nanoparticles (Fe_3_O_4_) to protect DNA and facilitate oligonucleotide transfer. Additionally, spermine-PCL-chitosan-polyethylene glycol (PEG)-folic acid (SPCPF) micelles were synthesized to control DNA release and target tumors. SCPPF/Fe_3_O_4_/DNA micelles were then prepared, and their properties, including protection against plasma degradation, ability to release DNA, physicochemical properties, transfection efficiency, and cytotoxicity, were evaluated *in vitro*. The study also investigated the impact of two buffer pH values (6 and 7.4) on DNA release from micelles, revealing that a lower pH significantly increased the release rate. Electrophoretic analysis confirmed that the micelle coating effectively protected DNA from degradation in plasma. VSM (vibrating sample magnetometer) analysis was used to assess the magnetic properties of the SCPPF/Fe_3_O_4_/DNA micelles, revealing that encapsulation with PCS and PCPF yielded nanoparticles with desirable magnetic characteristics. However, the encapsulation value reduced the saturation magnetic properties of the samples from 13.4 to 39.3 emu/g. Furthermore, the micelles significantly improved the DNA transfer efficiency compared with that of the polyethylenimine (PEI)/DNA complex in serum, achieving 14.42% efficiency with the SCPPF/Fe_3_O_4_/DNA micelles compared with 10.6% for the PEI/DNA complex. In terms of cytotoxicity, the SCPPF/Fe_3_O_4_ micelles exhibited low toxicity to the AGS cell line.

## Introduction

Cancer is a major global health challenge, causing significant mortality annually [1]. Metastasis complicates treatment, whereas the side effects of chemotherapy hinder effective management [2]. Gastric cancer, primarily adenocarcinoma, is a leading cause of cancer death in Asia. It results in approximately 1 million deaths worldwide each year [3].

Nanotechnology offers promising avenues for targeted therapies, reducing collateral damage to healthy tissues [4]. Gene therapy, which has advanced rapidly, is used to treat serious diseases such as cancer and genetic disorders [5]. While viruses are effective gene delivery tools, their potential for mutation poses challenges. Nanoparticles, such as liposomes, dendrimers, or polymeric micelles, provide safer and more efficient alternatives. They encapsulate genetic material, protect it from degradation, and facilitate transport through physiological barriers [6]. In response to stimuli such as pH changes or enzymes, nanoparticles release genes precisely, minimizing off-target effects and side effects [7]. Surface modifications with ligands or antibodies enable specific binding to overexpressed receptors on diseased cells, increasing transfection efficiency [8]. This ensures that genes reach their intended sites, correcting genetic disorders or promoting cancer cell apoptosis [9]. Nanoparticle-based gene therapy combines molecular biology and nanotechnology, improving therapeutic outcomes and offering a safer alternative to viral vectors [10].

Biodegradable polymers such as poly(lactic-co-glycolic acid) (PLGA), polylactic acid (PLA), and polycaprolactone (PCL) are highly biocompatible owing to their biodegradability and integration into the citric acid cycle [11]. The degradation rates of these materials, ranging from days to years, depend on the molecular weight, polymer type, and copolymer ratio [12]. Understanding these factors can help control DNA release rates in nanoparticles, especially for sustained-release applications [13]. However, PLA and PCL face challenges in gene delivery, such as large particle sizes, low DNA encapsulation efficiency, and hydrophobic interactions with plasma proteins, leading to rapid clearance by the reticuloendothelial system (RES) [14]. Recent research has explored zwitterionic segmental couplers, which combine hydrophilic and hydrophobic components, to improve drug delivery systems. Owing to their unique properties, these copolymers enhance the performance and effectiveness of drug delivery, making them valuable in this study.

Liposomes and polymers are key vectors for DNA delivery. Polyspermine, a natural polyamine with unique properties, has gained attention for targeted DNA delivery because of its biological compatibility and low toxicity. Its positive charge enables it to complex with negatively charged DNA, stabilizing it and facilitating delivery to cells [15]. Unlike traditional methods such as electroporation, which often cause cell damage and have low efficiency, polyspermine offers a safer and more effective alternative. It can be combined with liposomes or nanoparticles for precise gene delivery, minimizing off-target effects [16]. Additionally, polyspermine stabilizes enzymes involved in metabolic processes, enhances DNA transfer stability and efficiency, and supports DNA-dependent processes such as replication and gene expression. These features make polyspermine a promising tool for genetic therapy and cell biology research [17].

Chitosan, a cationic polysaccharide derived from chitin deacetylation, is widely used in drug delivery [18]. It dissolves in acidic solutions but is poorly soluble under neutral or basic conditions. Solubility depends on factors such as the degree of deacetylation, molecular weight, pH, temperature, and crystallinity, with greater deacetylation and a lower molecular weight enhancing solubility. Its positively charged amino groups enable mucoadhesive properties, facilitating interactions with negatively charged mucosal surfaces [19]. However, the poor mechanical properties and limited water solubility of chitosan necessitate modifications to its hydroxyl and amino groups to improve its effectiveness in tissue engineering and drug delivery applications [20]. Polycaprolactone (PCL), a hydrolytically degradable polymer synthesized by ring-opening polymerization of ε-caprolactone, is widely used in biological applications because of its controlled drug release profile, slow *in vivo* degradation, and ease of synthesis [21]. Its excellent mechanical properties and high drug permeability make it ideal for controlled drug delivery [22]. The chemical and physical properties of PCL can be tailored by modifying its monomers, enabling precise control over drug loading and release in PCL-based systems [23]. Polyethylene glycol (PEG), a hydrophilic polymer, is valuable in biomedical applications because of its nontoxicity, biocompatibility, nonimmunogenicity, and ability to reduce protein adsorption [24,25]. Its hydrophilicity can be adjusted by adjusting its molecular weight and temperature. With active hydroxyl ends, PEG effectively forms drug delivery systems with various pharmaceutical compounds, making it a versatile carrier material [26].

Recent advancements in drug delivery include functionalizing nanoparticle surfaces with folic acid (FA) to target cancer cells, particularly in gastric tumors, enhancing therapeutic efficacy while reducing systemic toxicity [27]. FA, a natural vitamin, improves biocompatibility and safety, and its receptors are overexpressed on many cancer cells, enabling precise targeting and increased drug uptake by cancer cells while sparing healthy cells [28].

Magnetic nanoparticles (MNPs), particularly iron oxide-based materials (Fe_3_O_4_), offer significant advantages for targeted drug delivery in treating gastric cancer (AGS) cells. Their ability to be guided by external magnetic fields ensures precise delivery to tumor sites, minimizing off-target effects and systemic toxicity [29]. MNPs exploit the enhanced permeability and retention (EPR) effect of tumors, allowing selective accumulation in cancerous tissues [30]. When functionalized with targeting ligands such as folic acid, they specifically bind to overexpressed receptors on AGS cells, enhancing their cellular uptake and therapeutic efficacy [15]. MNPs also enable controlled drug release in response to stimuli such as pH or magnetic fields, improving treatment precision. Additionally, they serve as multifunctional platforms for combined therapy, imaging, and hyperthermia, making them powerful tools for overcoming drug resistance and improving outcomes in gastric cancer treatment [31]. The biocompatibility, biodegradability, and ability of these materials to codeliver drugs and genes further underscore their potential in advanced cancer therapies. The choice of copolymer compounds in drug delivery systems is driven by their unique physicochemical properties, biocompatibility, biodegradability, and functionalities, which enhance drug solubility, targeted delivery, controlled release, and overall therapeutic efficacy [12]. By combining copolymers, researchers can tailor systems to meet specific therapeutic needs and improve patient outcomes.

In this study, we focused on DNA delivery into the cell nucleus, leveraging the high expression of FA receptors on cancer cell surfaces to enable targeted gene transfer to gastric cancer cells, minimizing off-target effects. We incorporated magnetic Fe_3_O_4_ nanoparticles with micellar copolymer nanoparticles to create a stable and homogeneous copolymer‒gene complex. Using the double emulsion solvent evaporation method, we synthesized Fe_3_O_4_-PCL-chitosan-PEG-SP-FA/DNA magnetic nanoparticles and evaluated their biocompatibility, DNA release rate, loading efficiency, and gene delivery efficacy in AGS gastric cancer cells.

## Materials and methods

The heterobifunctional PEG reagents with folic acid and amine groups (NH_2_-PEG-Folic acid) were purchased from Polysciences, Inc. (Germany) and Ruixi Biotechnology Co. (China), respectively. NH_2_-PEG-glucose and acrylate-PCL were prepared from Ruixibiotech (China). Dulbecco’s modified Eagle’s medium (DMEM) and trypsin-EDTA were purchased from Gibco (USA). Chitosan, chloroform, fetal bovine serum (FBS), penicillin/streptomycin solution, oleic acid, methyl sulfoxide (DMSO), 3-(4,5-dimethyl-2-thiazolyl)-2,5-diphenyl-2H-tetrazolium bromide (MTT) and RPMI 1640 were purchased from Sigma‒Aldrich (USA). Polyvinyl alcohol (PVA), Tris-HCl, agarose gel, FeCl_3_, FeCl_2_, and EDTA were purchased from Merck (Germany).

## Methods

### Synthesis of Spermine-Chitosan-PCL (PCS)

To synthesize PCS, we first prepared acrylate-chitosan-PCL. This involved dissolving 4 g of acrylate-PCL in 20 ml of chloroform, followed by the addition of 4 g of chitosan to the acrylate-PCL solution, which was stirred for 24 hours at 50°C. Afterward, methanol and water solvent exchange was performed to precipitate the product, which was subsequently dialyzed against water to remove impurities. The final mixture was stored at 4°C for 2 days to further eliminate contaminants [32]. To measure the concentration of chitosan-PCL, Fourier transform infrared (FTIR) spectroscopy was used. In this method, the concentrations of various components are determined by analysing the absorption peaks.

Next, chitosan-PCL (0.2 mmol) was reacted with 0.4 mmol of acetyl chloride in dry toluene containing 0.4 mmol of triethylamine. This mixture was vigorously stirred at 90°C for 6 hours. Once cooled to room temperature, the mixture was filtered to remove the triethylamine hydrochloride and was precipitated and purified via cold hexane. The resulting precipitate was then collected and vacuum-dried for 24 hours.

For the synthesis of spermine-chitosan-PCL, 2 g of acrylate-chitosan-PCL was dissolved in 20 ml of chloroform, to which 1 g of spermine was added. The mixture was stirred at 50°C for 24 hours, after which it was dialyzed to isolate the spermine-chitosan-PCL copolymer (**Fig. 1**).

**Fig. 1.**
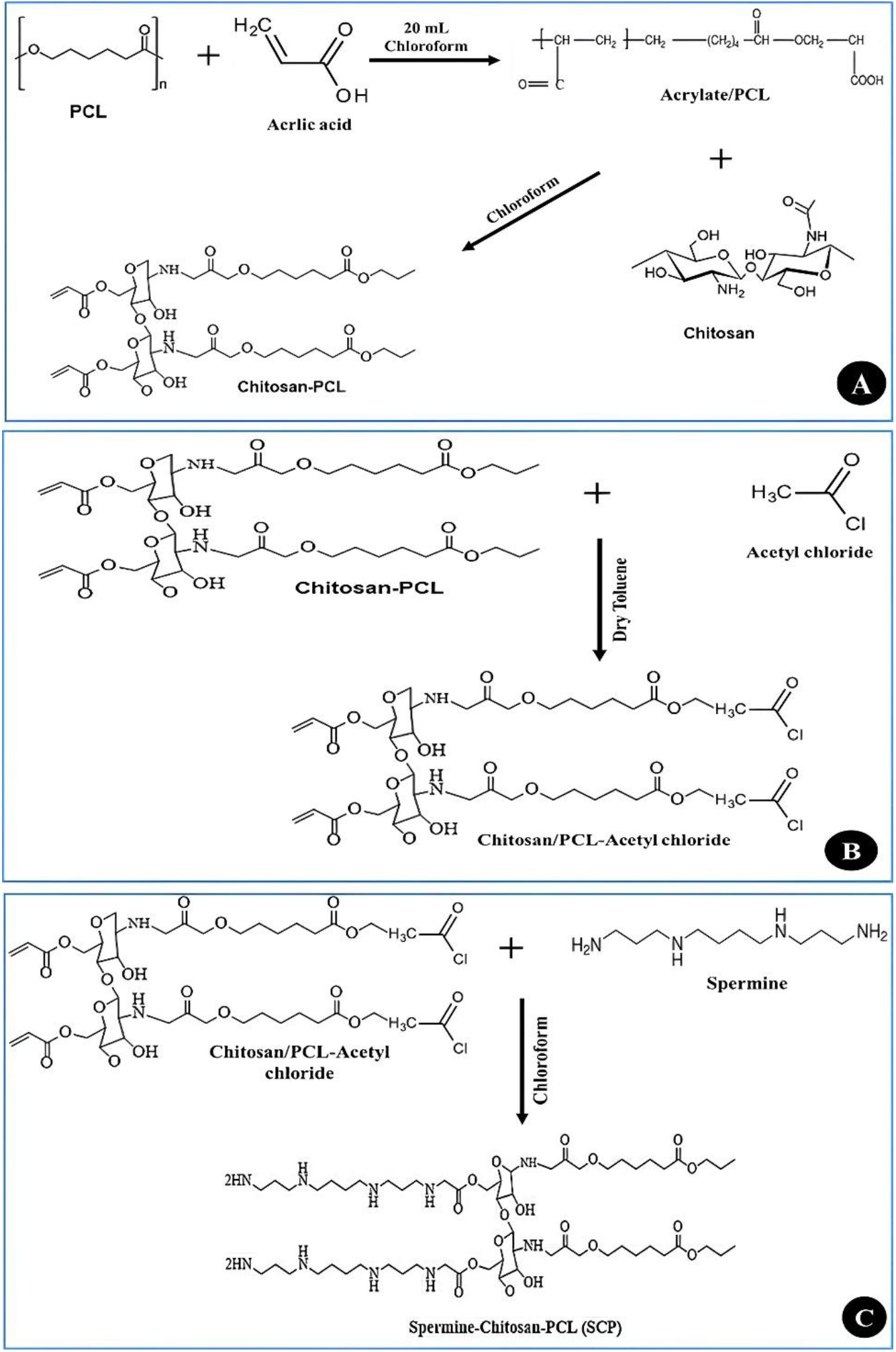
Chemical scheme of A) chitosan/PCL (CP) synthesis, B) chitosan/PCL-acetyl chloride synthesis, and C) spermine-chitosan/PCL (SCP) copolymer synthesis.

### Synthesis of Spermine-PCL-Chitosan-PEG-FA (SCPPF) and PCL-chitosan-PEG-FA (PCPF)

To synthesize spermine-PCL-chitosan-PEG-FA, we first dissolved 30 μmol of NH_2_-PEG-FA and 10 μmol of SCP in 3 ml of chloroform. The chloroform solution containing NH_2_-PEG-FA was then added dropwise to the chitosan-PCL solution, and the mixture was stirred at 40–45°C for 24 hours. Next, we purified the Spermine-PCL-Chitosan-PEG-FA solution through dialysis in deionized water for 2 days, changing the water every 6 hours. Finally, the SCPPF copolymer was obtained by freeze-drying (**Fig. 2**).

**Fig. 2.**
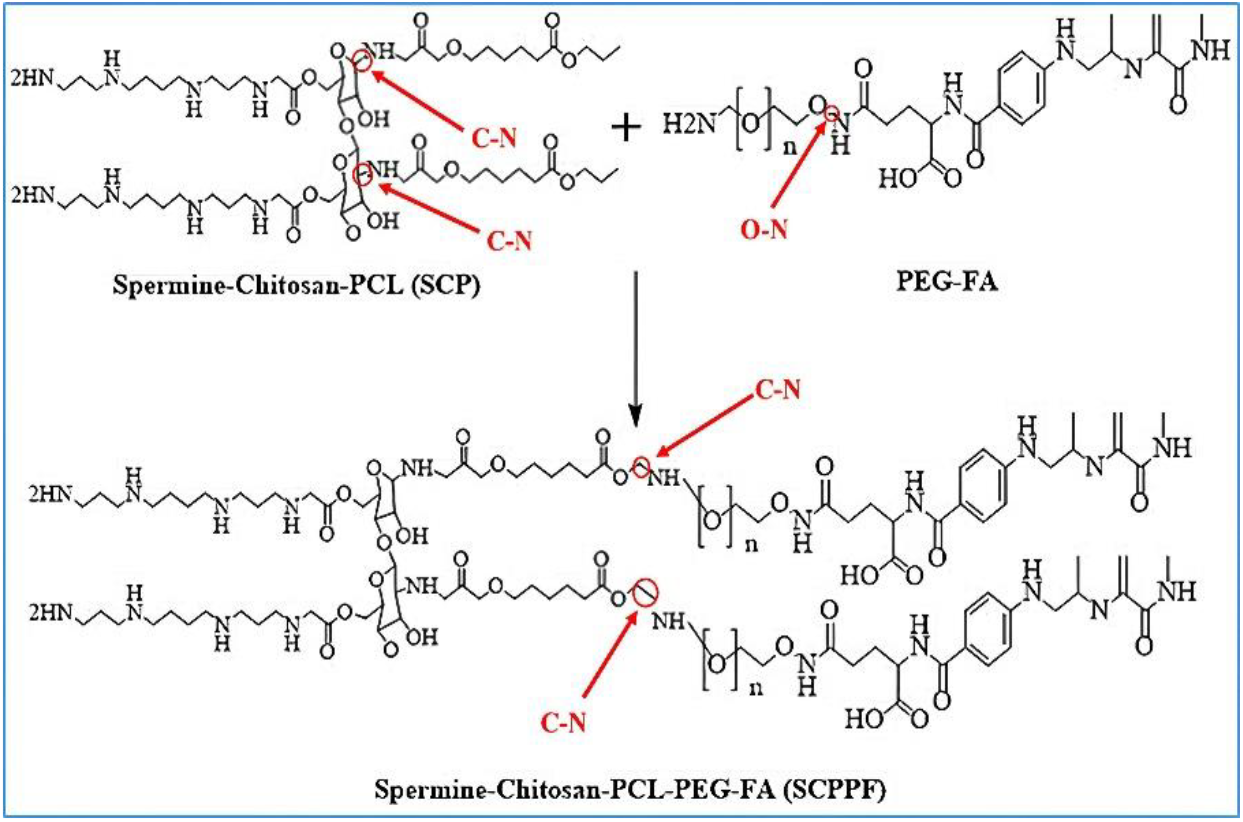
Chemical scheme of spermine-chitosan-PCL-PEG-FA (SCPPF) copolymer synthesis. In this figure, single covalent bonds are shown as C‒N and O‒N bonds.

### Green synthesis of iron nanoparticles (Fe_3_O_4_)

For the green synthesis of iron nanoparticles, we first prepared a hydroalcoholic extract using spinach leaves (*Spinacia oleracea* L.). The leaves were washed with deionized water, dried in the dark at room temperature, and then ground into a fine powder [33]. We combined 20 g of this powder with 100 ml of 80% methanol and incubated the mixture at 37°C with continuous shaking (50 rpm) for 24 hours. After incubation, we centrifuged the mixture at 16,600 × g for 10 minutes, removed the supernatant, and filtered it through filter paper.

Next, for the synthesis of Fe_3_O_4_, we mixed 3.33 g of FeCl_3_·6H_2_O and 1.59 g of FeCl_2_·4H_2_O with 100 mL of deionized water. This reaction was conducted under stirring with nitrogen gas at 80°C. We then added 15 ml of the spinach extract dropwise to the mixture and allowed it to react for 10 minutes. Next, we gradually introduced 60 ml of 100 mM sodium hydroxide (NaOH) solution until the color of the solution changed from brown to black. The resulting nanoparticles were collected via a strong neodymium magnet and dried after being rinsed with deionized water.

### Preparation of SCPPF/DNA micelles

Spermine-chitosan-PCL-Fe_3_O_4_-OA-PEG-FA/DNA (SCPPF/DNA) micelles were prepared via the double emulsion solvent evaporation method. Briefly, 250 μg of DNA (dissolved in 0.5 mL of TE buffer) was combined with 4 mL of a solvent mixture of acetone and DCM (50% each) that contained 10 mg of Fe_3_O_4_-oleic acid (OA) and 20 mg each of spermine-PCL-chitosan-PEG-FA and spermine-chitosan-PCL copolymers. This mixture was sonicated for 30 s at 0°C [34].

Next, we added 6 mL of a 1% (w/v) PVA solution to the emulsion and sonicated it for 2 minutes at 0°C. The emulsion was then added dropwise to 30 mL of a 0.3% (w/v) PVA solution while stirring mechanically for 30 minutes. After the solvents (acetone and DCM) were removed via a rotary vacuum evaporator (Heidolph, Germany), the nanoparticles were collected via centrifugation at 16602 × g for 30 minutes. Finally, the micelles were washed with deionized water and lyophilized through freeze-drying after filtration with 0.1- and 0.45-μm membrane filters to eliminate coarse particles and unbound copolymers.

To determine the efficiency of DNA encapsulation in micelles, the amount of DNA in the supernatant of each sample was measured via a Nanodrop (Thermo Scientific 2000, USA) at wavelengths of 260 and 280 nm and compared with the amount of DNA used in the samples.

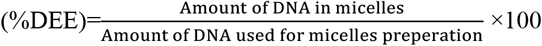

### Characterization of micelles

The synthesized copolymers were characterized by hydrogen nuclear magnetic resonance spectroscopy (^1^H-NMR-Bruker 400 MHz) and infrared spectroscopy (FTIR-ABB Bomem-MB). The morphology, particle size, mean diameter, polydispersity index (PDI), and zeta potential of the micelles were examined via transmission electron microscopy (TEM, JEM-2100), scanning electron microscopy (SEM), and dynamic light scattering (DLS, Malvern Instruments, Westborough, MA, USA), respectively.

The magnetic properties of the Fe_3_O_4_, Fe_3_O_4_-OA, and SCPPF/Fe_3_O_4_/DNA micelles were investigated via a vibrating sample magnetometer (VSM, Standard Series 7403, Lakeshore). The thermal properties of PCL, PEG, chitosan, spermine, PCS, and PCPF were determined via thermogravimetric analysis (TGA-DTA-32, Japan) at temperatures between 25°C and 600°C with a heating rate of 20°C/min under air pressure.

### Agarose gel electrophoresis of micelles

Agarose gel electrophoresis was conducted to assess the capacity of the micelles to encapsulate and neutralize the negative charge of the DNA. Following the preparation of micelles mixed with DNA, a gel was cast with 0.8% (w/v) agarose. Both the micelles and control DNA (with a constant amount of 2 μg of DNA per sample) were subjected to electrophoresis at a voltage of 80 V for 30 minutes [35].

### Cell culture

The AGS human gastric cancer cell line was obtained from the National Cell Bank of Iran (NCBI) at the Pasteur Institute (NCBI, C131). These cells were cultured in Dulbecco’s modified Eagle’s medium (DMEM) supplemented with 10% (v/v) heat-inactivated FBS and 1% penicillin‒streptomycin [36]. The cultures were maintained in a humidified incubator at 37°C with 5% CO_2_. To ensure continuous exponential growth, the cells were passaged weekly upon reaching 70%-80% confluence via the use of trypsin-EDTA.

### Preparation of PEI/DNA nanoparticles (N/P=5)

For this purpose, 25 kDa PEI was mixed with the psiCHECK plasmid. Then, 1 μg of DNA was mixed with different amounts of PEI on the basis of the calculation of N/P. The N/P ratio refers to the molecular ratio of amines (N; cationic groups) in PEI to phosphates (P; anionic groups) in nucleic acids. The N/P ratio was calculated from the molar ratio of positive to negative charge.

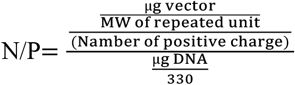

The molecular weight of a PEI repeating unit is 473 Da, with four amine groups:

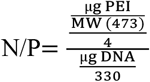

To obtain an N/P ratio of 5, 1 μg of DNA and 4.2 μg of PEI were mixed directly or diluted in 100 μl of PBS buffer. The mixtures were then incubated at 25°C for different durations of 5–30 min.

### *In vitro* cytotoxicity studies

The MTT assay is a widely employed method for evaluating cytotoxicity. This assay is based on the conversion of MTT into insoluble formazan crystals by viable cells, indicating mitochondrial activity [37]. As mitochondrial function typically ceases upon cell death, the MTT assay is a reliable measure of cell viability.

The cytotoxic effects of polyethylenimine (PEI)/DNA (N/P=5) and SCPPF/Fe_3_O_4_/DNA were assessed via the MTT assay. AGS cells were plated in 96-well plates at a density of 7 × 10^3^ cells per well in 200 μL of complete medium (RPMI 1640 supplemented with 10% FBS) and incubated at 37°C in a 5% CO_2_ atmosphere. After 24 hours, the cells were treated with various concentrations of PEI/DNA (N/P=5) and SCPPF/Fe_3_O_4_/DNA micelles for an additional 24 hours at 37°C. Cell viability was measured via the MTT assay at a wavelength of 570 nm (BioTek Instruments; Winooski, VT, USA) [38]. Finally, the percentage of surviving cells was determined via the following equation:

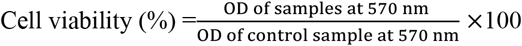

### DNA protection assay

To evaluate the protective capacity of micelles against plasma degradation, both micelles and uncoated DNA (with a fixed amount of 2 μg of DNA per sample) were incubated with 50 μL of 20% human plasma at 37°C for 30 minutes. Nucleases were inactivated by adding 5 μL of EDTA solution (0.5 M, pH=8), followed by the addition of heparin (1% w/v) and incubation at 37°C for 4 hours with shaking (Navarro & de ILarduya, 2009). The samples were subsequently analysed via gel electrophoresis via a 2% agarose gel at 80 V [39].

### Kinetics of the release of DNA from micelles

To assess DNA release, PEI/DNA and SCPPF/Fe_3_O_4_/DNA micelles were prepared as described earlier. Given the differing pH levels in normal and cancerous tissues, release assays were performed in two distinct media under acidic (pH=6) and neutral (pH=7.4) conditions. The PEI/DNA complex and micelles were incubated separately in 20 mL of PBS at 37°C. At scheduled intervals (ranging from 30 minutes to 30 days), the micelles were collected via centrifugation (16602 × g for 30 minutes), and the resulting supernatant was used for release assays. The micelles were then resuspended in fresh buffer and incubated for the next time interval. The amount of DNA in each sample was quantified via a NanoDrop spectrophotometer at a wavelength of 480 nm [40]. The cumulative percentage of DNA released from the nanoparticles was determined as follows:

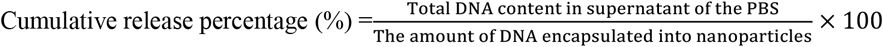

### Transfection assay

Transfection is a pivotal technique in molecular biology that enables the targeted introduction of genetic material into cells, facilitating the analysis of gene function, protein production, and cellular mechanisms [41]. This process involves delivering exogenous nucleic acids, such as DNA or RNA, into target cells. In this study, we evaluated the transfection efficiency of PEI/DNA complexes (with a nitrogen-to-phosphorus ratio of 5:1) and SCPPF/Fe_3_O_4_/DNA micelles in the AGS cancer cell line.

For the transfection assay, 1 ml of complete RPMI 1640 medium containing 2 × 10^4^ AGS cells was added to each well of a 24-well plate and incubated at 37°C with 5% CO_2_ for 24 h. After this incubation, the cells were washed twice with PBS buffer, and the culture medium was replaced with 1 ml of medium (with or without 10% FBS) containing 2 μg of DNA per micelle. The treatments included 2 μg of uncoated DNA (as a negative control), the PEI/DNA complex and the SCPPF/Fe_3_O_4_/DNA micelles. The cells were incubated under the same conditions for 8 h at 95% relative humidity. After 7 h, the transfection medium was replaced with fresh medium (1 ml), and the cells were incubated for another 48 h. DNA expression and the transfection efficiency of the micelles were assessed via an inverted fluorescence microscope (Nikon TE200) and flow cytometry (CyFlow Space, Germany) [42]. Lipofectamine 2000/DNA was also used as a positive control in this study.

### Statistical analysis

All the quantitative parameters examined in this study included a minimum of three replicates. Statistical analyses were performed via SPSS software, and one-way ANOVA was used to analyse the data. The Kolmogorov‒Smirnov test was used to assess the normality of the data set, whereas Duncan’s multiple-range test was applied at a 5% significance level for mean comparisons. The results for each treatment are presented as the means ± standard deviations (means ± SDsSDs).

## Results

### ^1^H-NMR results

^1^H-NMR spectroscopic examination of spermine-chitosan-PCL (SCP) and chitosan-PCL-PEG-FA (CPPF) revealed pronounced chemical shifts attributable to the hydrogen nuclei within their molecular frameworks (**Fig. 3A** and **B**). Chloroform is one of the most prevalent solvents employed in ^1^H-NMR spectroscopy. The utilization of chloroform in most spectrophotometric analyses is attributed to its consistent results and minimal interference with the analytes. Dimethyl sulfoxide (DMSO) is an additional solvent extensively utilized across various spectroscopic techniques and is recognized for its polar characteristics, which facilitate the dissolution of a multitude of organic and biological substances. A significant advantage of employing dual solvents in ^1^H-NMR spectroscopy is the ability to differentiate analogous signals effectively. Furthermore, the concurrent use of dual solvents enables a comprehensive examination of the influence of solubility on the activation energies and spectroscopic peaks. Consequently, in the present investigation, we employed two polar solvents, namely, chloroform and DMSO, as the media for the nanoparticles within the framework of ^1^H-NMR spectroscopy. The peaks discerned from chitosan in the ^1^H-NMR SCP spectrum are delineated as follows: the peaks corresponding to the methyl groups of chitosan are detected in the 4 to 5 ppm range in ^1^HNMR SCP spectroscopy (**Indexes 6, 7, 8, and 10 in Fig. 3 A**).

**Fig. 3.**
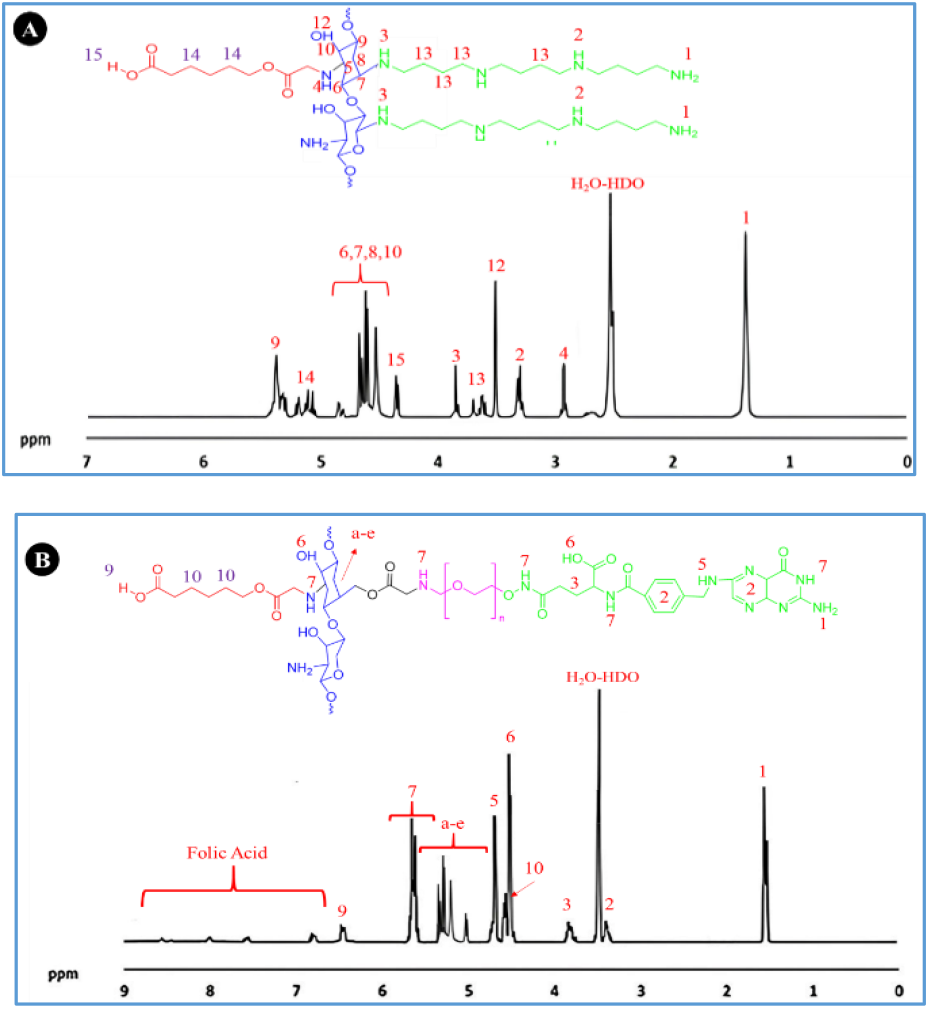
^1^H-NMR spectra of (A) Spermine-chitosan-PCL (SCP) and (B) Chitosan-PCL-PEG-FA (CPPF).

The peak corresponding to methylene groups was detected between 5.3 and 5.5 ppm. These protons are situated near hydroxyl and amine groups, and their chemical shifts are influenced by the electronegativity of the oxygen and nitrogen atoms. Additionally, regarding the nitrogen (NH) peaks, chitosan contains nitrogen from amino groups alongside hydrogen and carbon. The NH proton peaks typically appear in the 2 to 3 ppm range, indicating the presence of amino groups in chitosan. These peaks are valuable for analysing chemical reactions and structural alterations. The peak at approximately 2.9 ppm can be associated with the proton of the N-H group (**Index 4 in Fig. 3 A**). Furthermore, the strong peaks observed at approximately 1250 ppm indicate the presence of the 2NH group in the PLA polymer, as observed via ^1^H-NMR spectroscopy of the SCP copolymer (**Index 1 in Fig. 3A**). The peaks between 3 and 4 ppm are attributed to CH_3_ group stretching in the PLA polymer (**Index 13 in Fig. 3 A**). The NH protons in the PLA polymer also appear in the 3 to 4 ppm range (**Indexes 3 and 4 in Fig. 3 A and B**), with peak variations attributed to the high electronegativity of oxygen. Additionally, peaks associated with the O-H group in polyspermine were found at approximately 4.4 ppm (**Index 15 in Fig. 3 A and B**). After the attachment of PEG-FA to SCP, new peaks emerged in the 7 to 9 ppm range, corresponding to the folic acid group.

### FTIR results

The results of the FTIR spectroscopy of chitosan revealed that the peaks in the ranges of 1500–1600 cm^−1^ and 3200-3500 cm^−1^ were related to N–H and O–H, respectively. Additionally, stretching related to the C-O functional group in the chitosan structure was observed in the range of 1020-1150 cm^−1^ (**Fig. 4 A**). Additionally, according to the results of the FTIR spectroscopy of PCL (**Fig. 4 B**), the peaks in the ranges of 1000-1500 cm^−1^ and 1500-2000 cm^−1^ were related to C–O and C=O, respectively (**Fig. 4 B**).

**Fig. 4.**
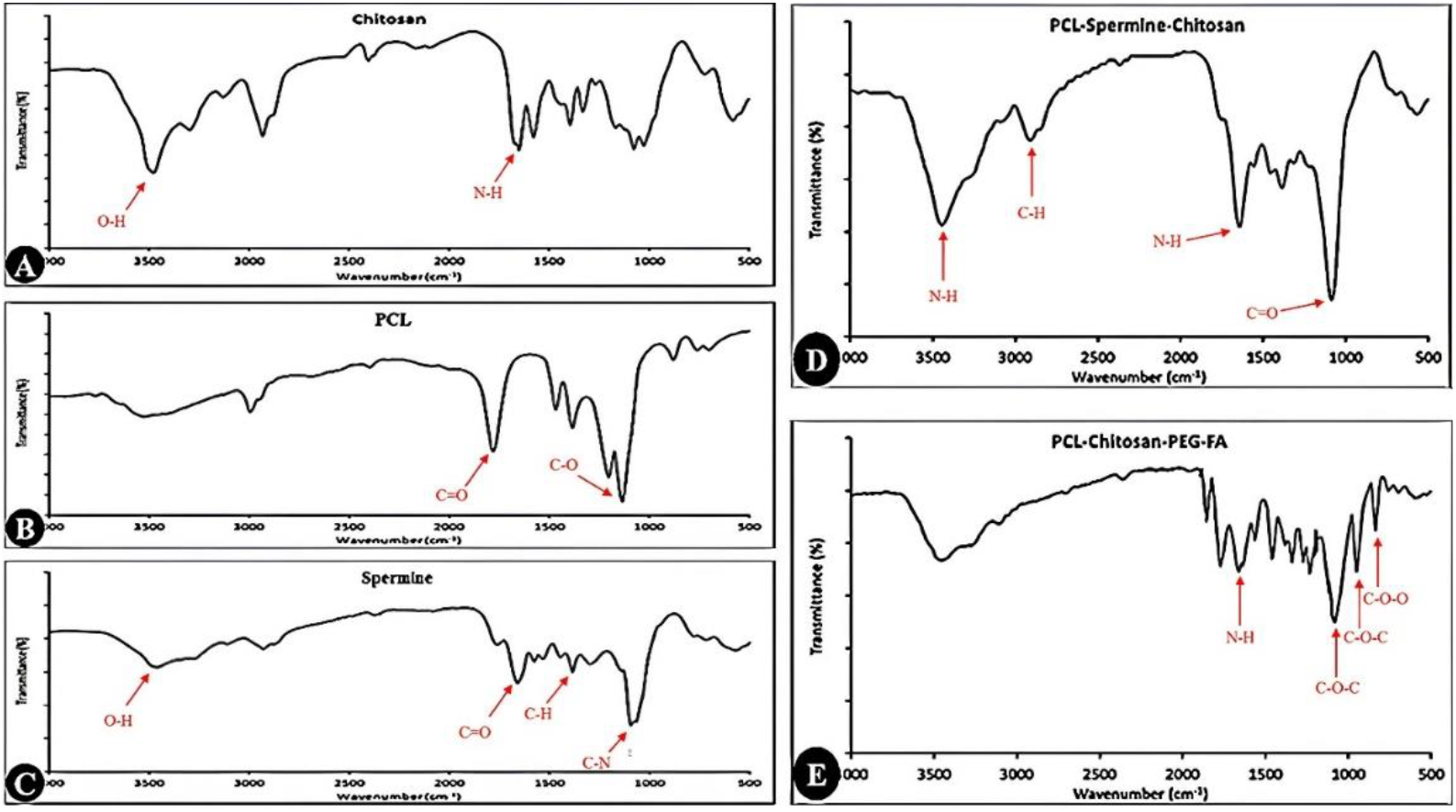
FTIR spectra of (A) chitosan, (B) PCL, (C) spermine, (D) spermine-chitosan-PCL (PCS), and (E) chitosan-PCL-PEG-FA.

The results of the FTIR spectroscopy of spermine showed that peaks related to O-H functional groups were observed in the range of 3360-3500 cm^−1^. The peak observed in the region of 1100 cm^−1^ corresponded to the CH_2_ group. Additionally, stretching related to the C‒N functional groups in the spermine structure was observed in the range of 1073 cm^−1^ (**Fig. 4 C**).

The FTIR spectrum of PCL-chitosan-spermine displays key bands at 1163 cm^−1^ and in the range of 2500-3000 cm^−1^, corresponding to the C=O stretching and C-H groups of PCL, respectively. The strong peak at 1559 cm^−1^ corresponds to the N‒H deformation of amide bond II in chitosan. Additionally, a peak at 3380 cm^−1^ is attributed to the primary amines of spermine (**Fig. 4 D**). For the PCL-chitosan-PEG-FA copolymers, peaks at approximately 948 cm^−1^ and 1348 cm^−1^ indicated C-O-C stretching related to PEG, confirming its grafting onto these copolymers. Weak peaks in the 1580-1650 cm^−1^ region indicated the presence of amino groups from folic acid (**Fig. 4 E**).

### Thermogravimetric analysis (TGA)

**Fig. 5 A and B** present the TGA profiles of PCL, chitosan, PEG-FA, PCL-chitosan-spermine, and PCL-chitosan-PEG-FA. Generally, all the compounds experienced weight loss ranging from 2% to 25% as the temperature increased from zero to 200°C, primarily due to the evaporation of residual water in the samples. As illustrated in **Fig. 5 A**, there were significant differences in thermal stability based on the material composition. PCL presented the highest thermal resistance, whereas chitosan presented the lowest. Our results indicated that, compared with both PEG-FA and PCL, spermine had greater thermal resistance.

**Fig. 5.**
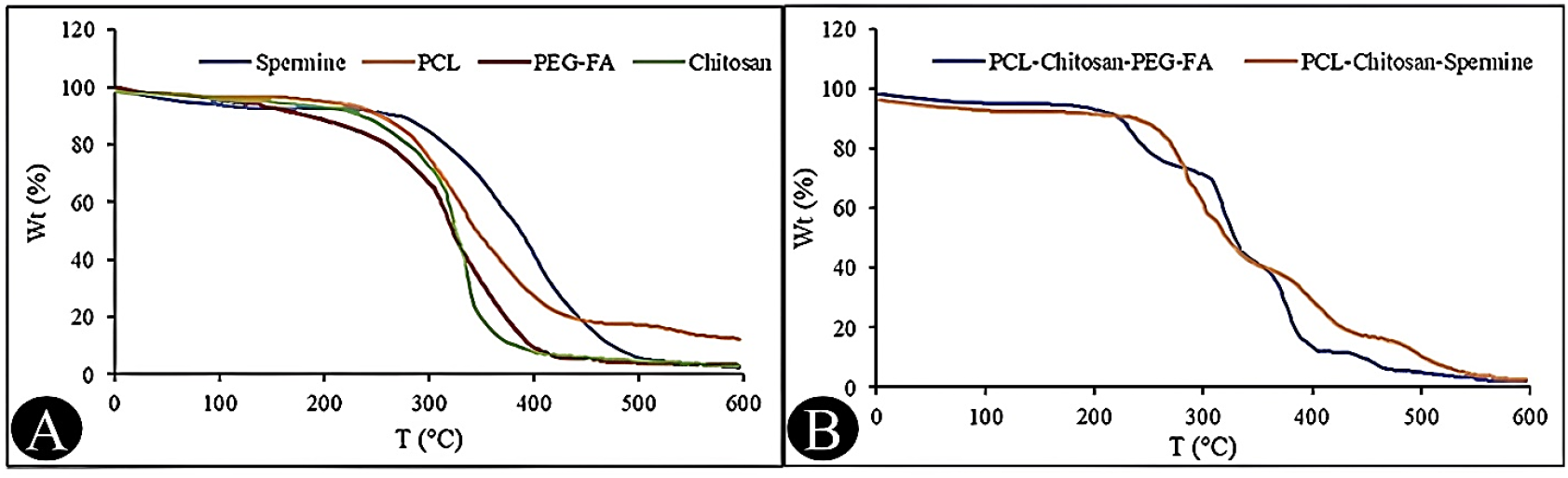
TGA analysis of (A) spermine, PCL, PEG-FA, and chitosan and (B) PCL-chitosan-PEG-FA and PCL-chitosan-spermine

The tricopolymer PCL-chitosan-spermine increased the number of weight loss steps. For example, the TGA curve for PCL demonstrated three distinct weight loss stages at temperatures ranging from zero to 200°C, 250 to 350°C, and 550 to 600°C. In contrast, the inclusion of PEG-FA increased the number of weight loss stages to more than five. A similar trend was observed for PCL-chitosan-spermine (**Fig. 5 B**).

### Physicochemical characterization of SCPPF/Fe_3_O_4_/DNA micelles and Fe_3_O_4_ nanoparticles

Controlling the morphology of nanoparticles is crucial for optimizing their transfection efficiency and drug release kinetics [43]. The transmission electron microscopy (TEM) results for the Fe_3_O_4_, Fe_3_O_4_-OA, and SCPPF/Fe_3_O_4_/DNA micelles revealed that the Fe_3_O_4_ nanoparticles exhibited a spherical shape with diameters ranging from 25–40 nm (**Fig. 6**). Additionally, both the TEM and SEM images indicated that the SCPPF/Fe_3_O_4_/DNA micelles were also spherical, characterized by a smooth surface and uniform distribution (**Fig. 6**). This spherical morphology of the micelles typically arises from the self-assembly of amphiphilic block copolymers in aqueous solutions.

**Fig. 6.**
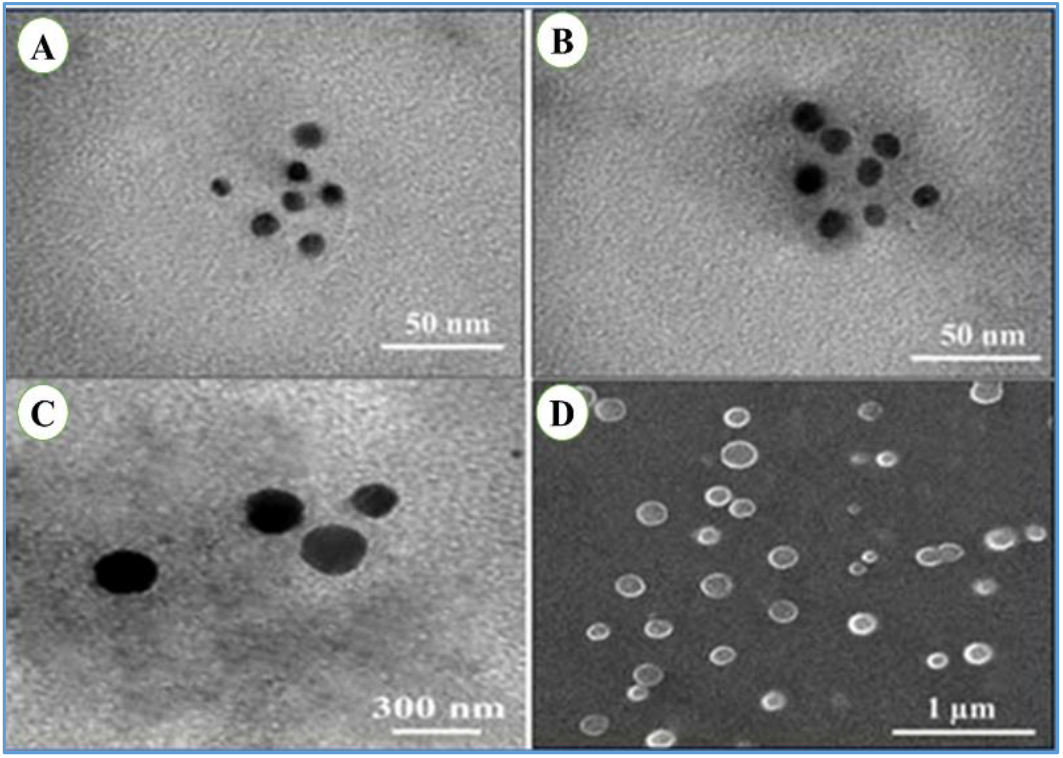
Morphology of the investigated nanoparticles: TEM images of the Fe_3_O_4_, Fe_3_O_4_-OA, and SCPPF/Fe_3_O_4_/DNA micelles (A, B and C), SEM image of the SCPPF/Fe_3_O_4_/DNA micelles (D).

The DLS plots for the Fe_3_O_4_ nanoparticles and SCPPF/Fe_3_O_4_/DNA micelles revealed a peak with a suitable amplitude, indicating uniformity among the nanoparticles (**Fig. 7 A and B**). The size of the Fe_3_O_4_ nanoparticles was approximately 39 nm, whereas the size of the SCPPF/Fe_3_O_4_/DNA micelles was approximately 265 nm (**Table 1**). The larger size of the micelles reveals successful polymer binding. The sharp peaks in the DLS plots of both types of nanoparticles (**Fig. 7 A and B**) indicated a uniform size, which is desirable for consistent biological behavior and effective drug delivery.

**Table 1.** Size and zeta potential of the Fe_3_O_4_ and SCPPF/Fe_3_O_4_/DNA micelles.

|  | Size (nm) | Standard deviation | Zeta potential (mV) | Standard deviation | PDI | Standard deviation |
| --- | --- | --- | --- | --- | --- | --- |
| $Fe_3O_4$ | 39 | $\pm 1.7$ | +1.2 | $\pm 0.09$ | 0.17 | $\pm 0.009$ |
| SCPPF/$Fe_3O_4$/DNA | 265 | $\pm 2.1$ | -2.5 | $\pm 0.12$ | 0.51 | $\pm 0.02$ |

**Fig. 7.**
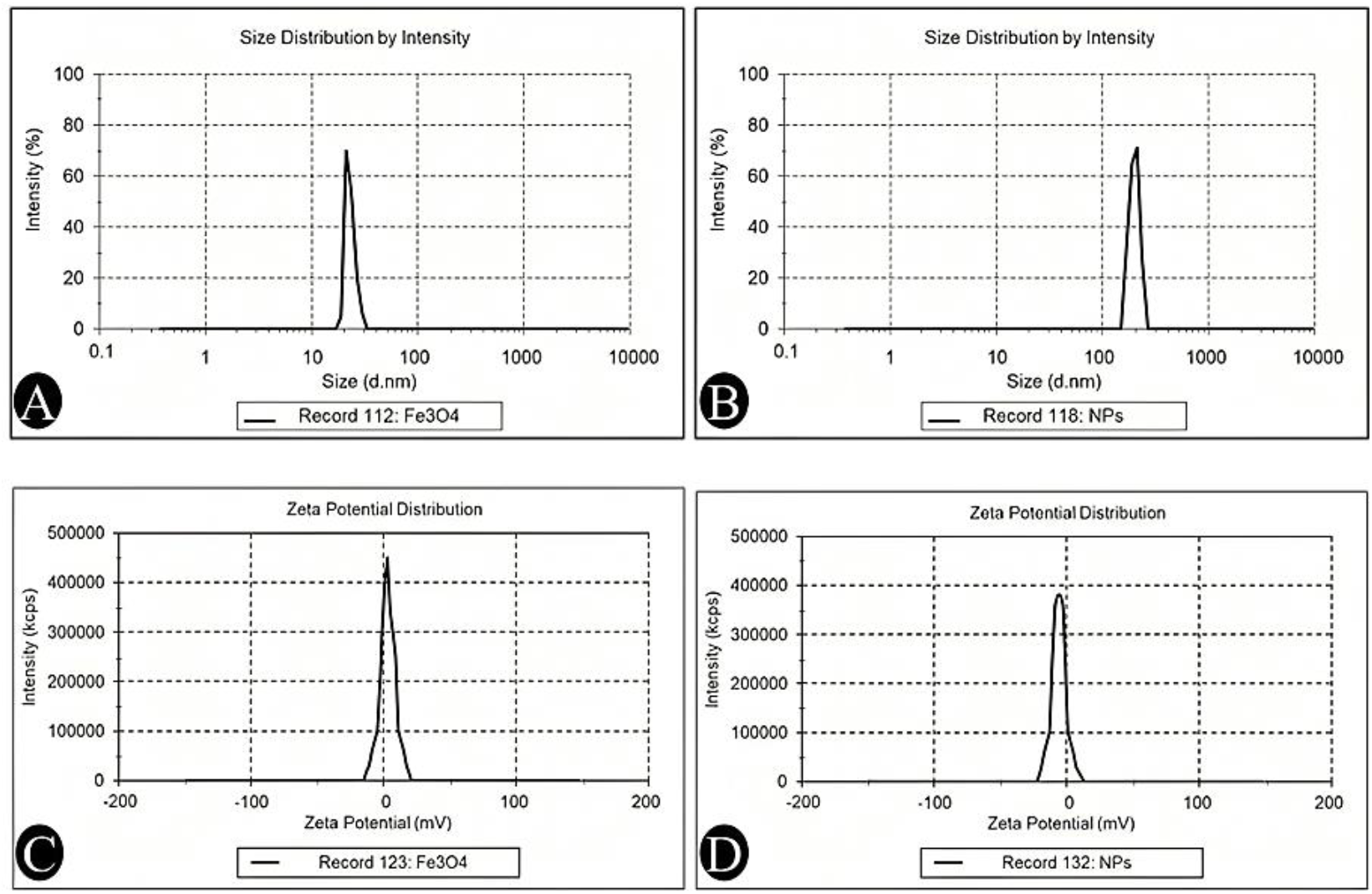
DLS and zeta potential diagrams of A, B) size distributions of Fe_3_O_4_ and SCPPF/Fe_3_O_4_/DNA micelles, respectively, and C, D) zeta potentials of Fe_3_O_4_ and SCPPF/Fe_3_O_4_/DNA micelles, respectively.

PDI is a measure of the distribution of molecular mass in a given sample. In the context of nanoparticles, the PDI provides insight into the uniformity of the particle size. A lower PDI indicates a more uniform sample, whereas a higher PDI suggests a broader size distribution. The PDI values of the nanoparticles were determined as follows:

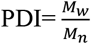

(M_w_= weight average molecular weight, M_n_= number average molecular weight) The following formulas were used to calculate the M_W_ and M_n_ values: where N_i_ is the number of particles and M_i_ represents the mass of the particles.

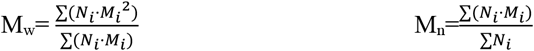

The mean diameter and PDI are fundamental parameters since they determine the *in vivo* transport of nanoparticles. The PDI value can range from 0.01 (monodispersal particles) to 0.5-0.7, whereas the broad particle size distribution of the formulation was indicated by a PDI index value > 0.7. In this study, the PDI value of the Fe_3_O_4_ nanoparticles was 0.17, indicating their homogeneity. However, after the synthesis and encapsulation of the SCPPF/Fe_3_O_4_/DNA micelles, the PDI value increased to 0.51, which indicates the successful binding of the spermine, chitosan, PLA, PEG, and FA copolymers (**Table 1**). Overall, the preparation method used in this study yielded nanoparticles with appropriate sizes and distributions.

The zeta potential distribution reflected the electrostatic stability of the nanoparticles. The Fe_3_O_4_ nanoparticles displayed an average positive charge of +1.2 mV (**Table 1 and Fig. 7 C**), whereas the SCPPF/Fe_3_O_4_/DNA micelles presented a negative zeta potential of −2.5 mV (**Table 1 and Fig. 7 D**). The decrease in zeta potential may result from the nanoparticles being surrounded by folic acid, and the presence of DNA sequences likely contributes to the greater negative charge due to their low surface charge [44].

### Magnetic properties of the micelles

Owing to their strong magnetic properties, Fe_3_O_4_ nanoparticles have significant applications in targeting and imaging systems [45]. The saturation magnetic properties of the samples studied ranged from 13.4 to 39.3 emu/g (electromagnetic units per gram). The highest magnetic properties were observed for the Fe_3_O_4_ iron oxide nanoparticles, with a value of 39.3 emu/g, whereas this value was 28.5 emu/g for the Fe_3_O_4_-OA nanoparticles. After encapsulating the Fe_3_O_4_-OA nanoparticles with PCL-chitosan-spermine and PCL-chitosan-PEG-FA micelles, the magnetic properties of the SCPPF/Fe_3_O_4_/DNA nanoparticles decreased to 13.4 emu/g. Overall, the saturation magnetic properties of the micelles diminished with increasing PCS concentration (**Fig. 8**). Nevertheless, the values reported in **Fig. 8** suggest that these results are suitable for biomedical applications.

**Fig. 8.**
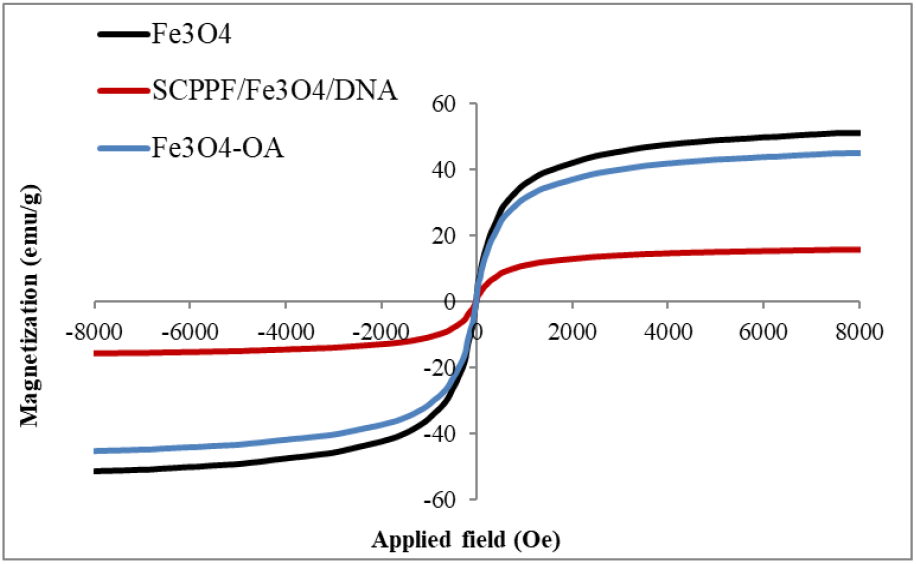
Magnetic properties of the Fe_3_O_4_, Fe_3_O_4_-OA, and SCPPF/Fe_3_O_4_/DNA micelles.

### Investigating the percentage of DNA release pattern from SCPPF/Fe_3_O_4_/DNA micelles

The DNA loading capacity of the SCPPF/Fe_3_O_4_/DNA micelles was 32.8%. The investigation into the DNA release pattern from SCPPF/Fe_3_O_4_/DNA micelles under neutral (pH=7.4) and acidic (pH=6) conditions revealed that, in general, DNA release from the PEI/DNA complex occurs more rapidly in a pH=7.4 environment with high concentrations of dextran than in SCPPF/Fe_3_O_4_/DNA nanoparticles. At a dextran concentration of 50 mg/ml, the percentages of DNA released from the PEI/DNA complex and the SCPPF/Fe_3_O_4_/DNA micelles were 42.3% and 18.64%, respectively. In contrast, the number of DNA strands in the acidic environment was significantly greater than that in the neutral environment. Additionally, no significant difference was observed in the amount of DNA released from the PEI/DNA complex or the SCPPF/Fe_3_O_4_/DNA micelles (**Fig. 9**).

**Fig. 9.**
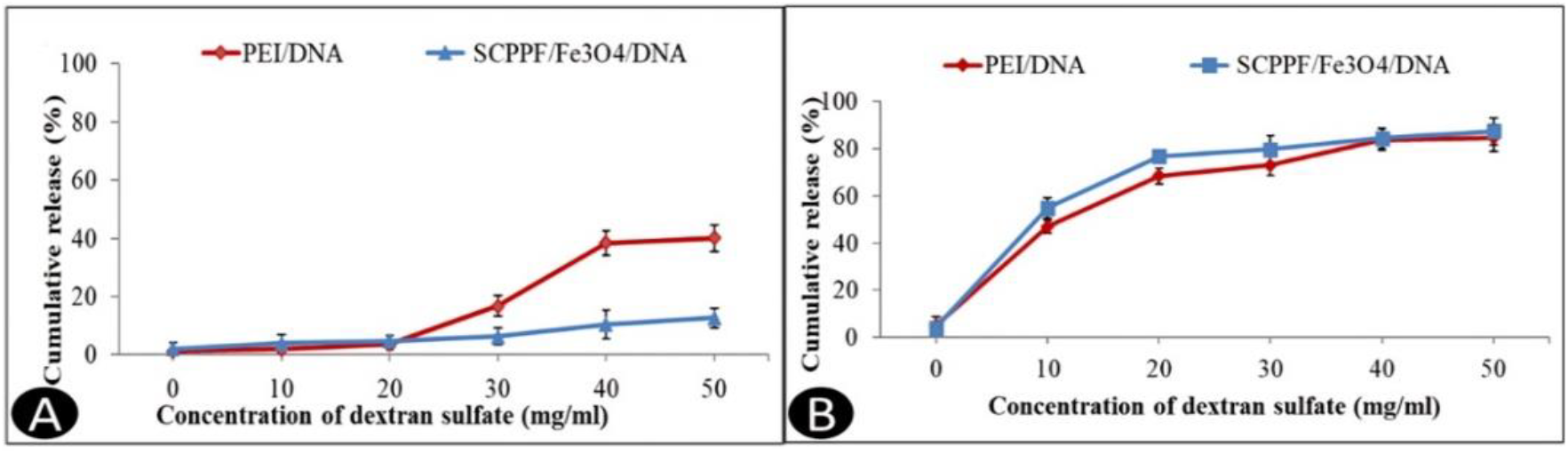
Drug release patterns of PEI/DNA and SCPPF/Fe_3_O_4_/DNA micelles in the presence of dextran sulfate at different pH values: A) pH=7.4 and B) pH=6.

### *In vitro* cytotoxicity assay

According to the results obtained from the analysis of variance table (**Table 2**), the viability of the AGS cell line was significantly (p < 0.01) affected by the type and different concentrations of the treatments used, such as Fe_3_O_4_, PCL-chitosan/PEG-FA (CPPF), PEI, SCPPF/Fe3O4/DNA, PEI/DNA and quercetin.

**Table 2.** Analysis of variance table of viability of AGS cell lines under the influence of different treatments.

| Sources of Variation | df | Mean Squares |
| --- | --- | --- |
|  |  | Cell viability (%) |
| Treatment | 29 | 1598.60** |
| Error | 60 | 1.41 |
| Coefficient of Variation (%) (CV) | - | 15.89 |

\*\* : Significant at the one percent probability level, and df: degrees of freedom.

One of the most important factors for the use of nanoparticles in clinical applications is their biocompatibility [7]. According to the results presented in **Fig. 10**, the Fe_3_O_4_ nanoparticles demonstrated very low toxicity at concentrations less than 200 μg/mL. However, increasing the concentration of this compound to 400 μg/mL negatively affected the viability of AGS cells, although this effect was not significant compared with that at concentrations of 100 and 200 μg/mL. Similar results were observed with PCL-chitosan/PEG-FA (CPPF) treatment. Furthermore, the results showed that the cytotoxicity of PEI (with or without DNA) was significantly greater than that of the positive control treatment, quercetin (Que), which led to a decrease in cell viability and death of up to 68.78% of AGS cells. The lowest cell viability (19.89%) was also recorded in the cell sample treated with 400 μg/mL quercetin. However, as shown in **Fig. 10**, the SCPPF/Fe_3_O_4_/DNA nanoparticles had lower cytotoxicity, with the viability of AGS cells treated with the highest concentration, i.e., 400 μg/mL SCPPF/Fe_3_O_4_/DNA nanoparticles, being 80.67%. Considering this, it can be concluded that the use of SCPPF/Fe_3_O_4_/DNA nanoparticles results in lower toxicity to cells and is very beneficial for the targeted delivery of genomic sequences while having minimal impact on the host cell.

**Fig. 10.**
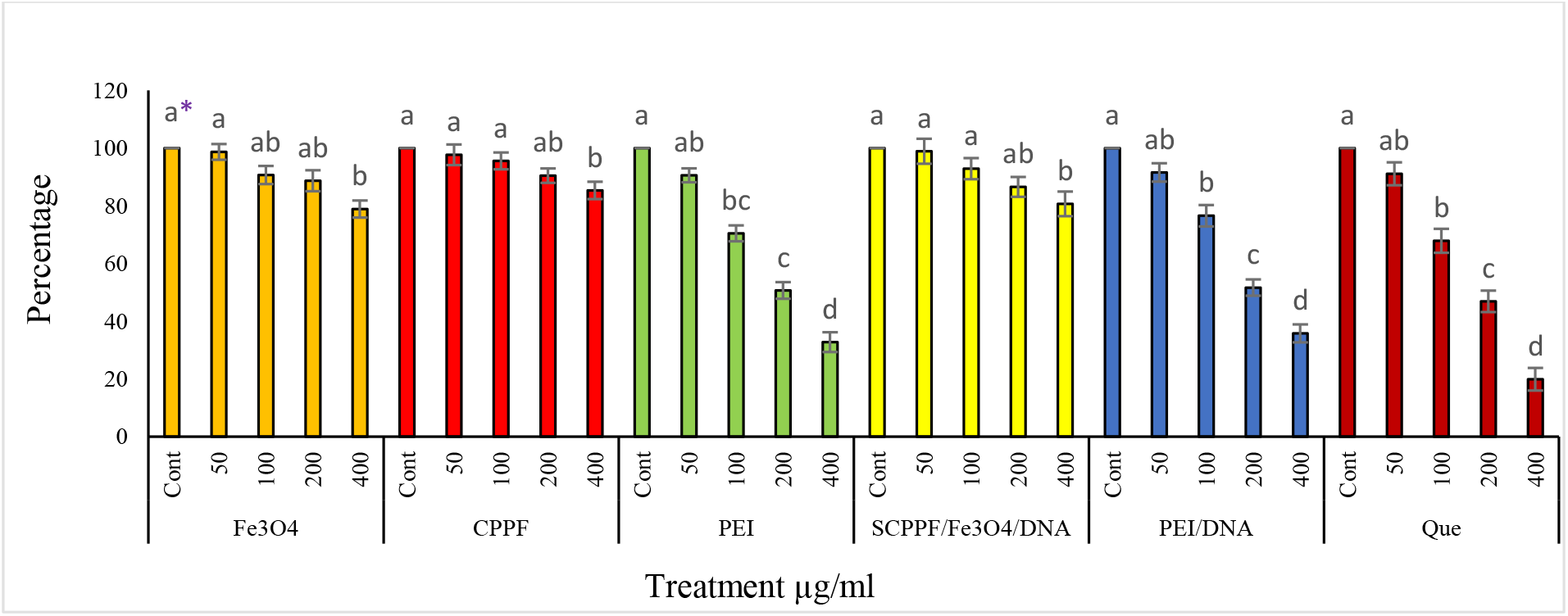
Cytotoxicity of the AGS cell lines treated with Fe_3_O_4_, chitosan-PCL/PEG-FA/(CPPF), PEI, spermine-PCL-chitosan/PEG-FA (SCCPF)/Fe_3_O_4_/DNA, PEI/DNA, and quercetin (Que).

^*^Different letters indicate significant differences (at the 5% probability level) among the treatments at each of the studied time points.

### Protection of DNA by Micelles from Degradation

The interaction of micelles with DNA and their protective effects against nuclease enzymes were examined through agarose gel electrophoresis [6]. As shown in **Fig. 11**, the SCPPF/Fe_3_O_4_/DNA nanoparticles effectively interact with DNA. By neutralizing the negative charge of DNA, these nanoparticles prevent movement through the agarose gel, causing the DNA in the well to remain immobile. Additionally, the DNA isolated from the micelles remained intact after incubation with human plasma (**Fig. 11**). In contrast, when the uncoated DNA was treated with human plasma, no visible DNA band appeared on the agarose gel, indicating degradation. This degradation results in the formation of small DNA fragments, which migrate through the agarose gel more quickly. These results suggest that the SCPPF/Fe_3_O_4_/DNA nanoparticles successfully protected the encapsulated DNA from nuclease digestion.

**Fig. 11.**
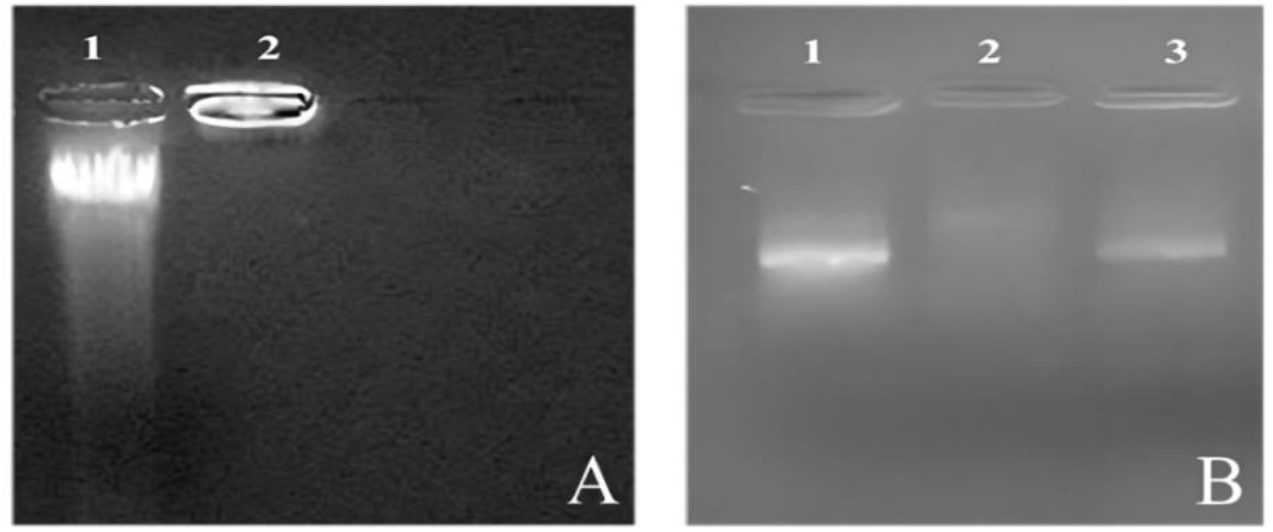
Effect of SCPPF/Fe_3_O_4_/DNA micelles on the interaction (A) with DNA. First well: control DNA; second well: SCPPF/Fe_3_O_4_/DNA nanoparticles; (B) Effect of SCPPF/Fe_3_O_4_/DNA micelles on protecting DNA against enzymatic digestion. First well: control DNA without plasma treatment; second well: control DNA treated with plasma; and third well: DNA extracted from SCPPF/Fe_3_O_4_/DNA micelles after plasma treatment.

### Cellular Transfection Assay

Transfection is a method used to introduce foreign nucleic acids into eukaryotic cells, altering the genetic composition of the host cell [9]. By understanding the molecular pathways associated with diseases, specific biomarkers that may aid in diagnosing and predicting outcomes can be identified. Additionally, transfection can be utilized as a strategy in gene therapy to address incurable inherited genetic disorders. The ability of SCPPF/Fe_3_O_4_/DNA nanoparticles to deliver DNA into AGS cells was examined via flow cytometry and fluorescence microscopy. Our findings (**Fig. 12**) clearly illustrate the effectiveness of these nanoparticles in delivering and releasing DNA into AGS cells. The transfection efficiency of PEI/DNA was 10.6%, whereas that of the SCPPF/Fe_3_O_4_/DNA nanoparticles reached 42.14% (**Fig. 13**).

**Fig. 12.**
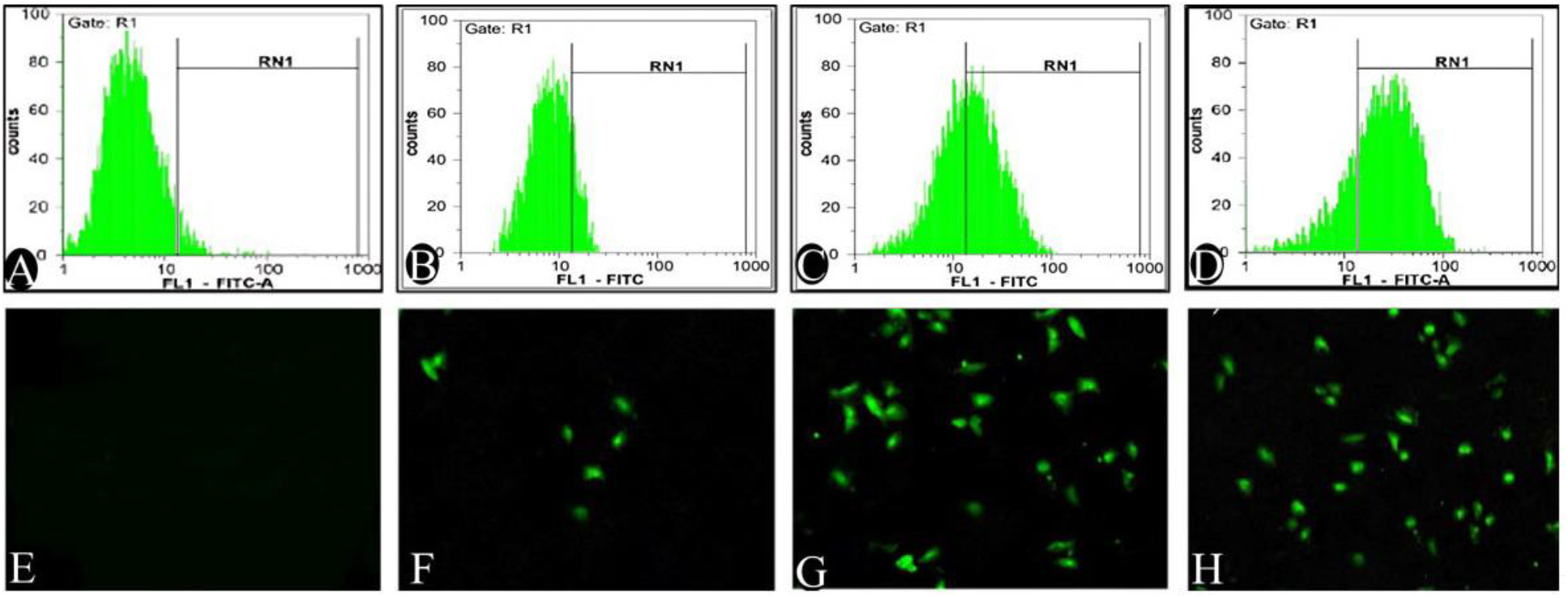
Transfection efficiency results and flow cytometry plots of treated AGS cells. A-D) Flow cytometry results of MCF-7 cells incubated with naked DNA (negative control), PEI/DNA, SCPPF/Fe_3_O_4_/DNA and Lipofectamine 2000/DNA (positive control). E-H) Fluorescence microscopy images of MCF-7 cells incubated with naked DNA (negative control), PEI/DNA, SCPPF/Fe_3_O_4_/DNA and Lipofectamine 2000/DNA (positive control).

**Fig. 13.**
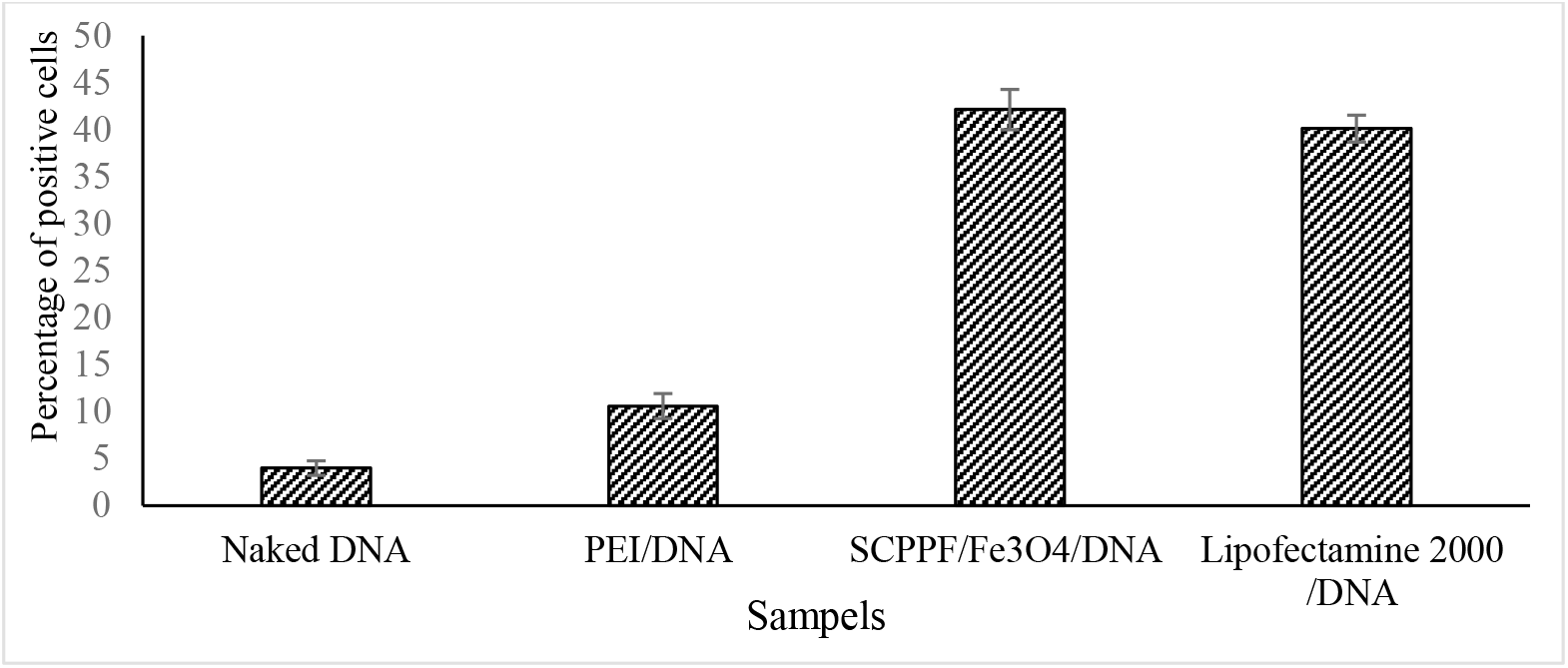
Comparison of the average gene transfer efficiency of the SCPPF/Fe_3_O_4_/DNA nanoparticles to MCF-7 cells

## Discussion

Recent advancements in disease diagnosis and treatment have enabled the management of many conditions, yet effective treatments for numerous genetic diseases remain elusive. Gene therapy offers a solution by correcting damaged genes or suppressing harmful gene expression [46]. However, delivering uncoated DNA to cells is challenging because of its negative charge, large size, and susceptibility to degradation [47]. Cationic nanoparticles can neutralize this negative charge through electrostatic interactions with DNA, facilitating DNA packaging, endosomal escape via the proton sponge effect, and entry into the cytoplasm or nucleus [48]. PEI is an efficient gene carrier, but its high toxicity to cancer cells limits its clinical use. Research has explored the toxicity mechanisms of cationic polymers such as PEI, revealing that the strong electrostatic interactions between the amine groups of PEI and cell surface phosphates reduce cell viability by disrupting the cell membrane and creating pores [49]. To investigate the toxicity of PEI, various strategies, including covalent bonding with biocompatible polymers, have been employed.

In this study, a complex of multiple nanoparticles was utilized for successful and targeted DNA transfer to AGS cells, taking into account the benefits and drawbacks of each individual nanoparticle. Biocompatible substances such as chitosan, PCL, PEG, spermine, and folic acid have drawn much interest in the field of nanomedicine, especially because of their potential applications in gene delivery systems [50]. All these elements possess distinct characteristics that offer benefits and drawbacks when used separately. Combining these methods, however, might improve therapeutic results and the effectiveness of gene delivery. Because of its positive charge at physiological pH, chitosan, a naturally occurring polymer derived from chitin, is known for its biocompatibility, biodegradability, and capacity to promote cellular uptake. Owing to their mucoadhesive qualities, nanoparticles are better retained in biological systems, which raises the possibility of gene transfer. Nevertheless, the stability and encapsulation effectiveness of chitosan-based nanoparticles may be impacted by their limited mechanical strength. PCL is a synthetic polyester that has low toxicity and good biodegradability. It is a good option for prolonged drug release because of its slow rate of degradation. Although PCL can create nanoparticles that shield genetic material from deterioration, occasionally, this material lacks bioactivity, which can impede its cellular uptake and interaction.

The hydrophilicity and ability of PEG to produce a covert effect, which reduces immune system recognition and clearance, make it a popular choice for drug delivery systems. Although excessive PEGylation can shorten the time of nanoparticle circulation in the bloodstream, it can also decrease the ability of nanoparticles to interact with target cells, which is important for efficient gene delivery. A naturally occurring polyamine called spermine can greatly increase the cellular uptake of nanoparticles by promoting endosomal escape. It is important to carefully optimize formulations because the efficacy of spermine can vary on the basis of formulation and concentration [51]. The affinity of folic acid for folate receptors, which are frequently overexpressed in different cancer cells, makes it a targeting ligand. To use this targeting capability, which improves the specificity and effectiveness of gene delivery, the nanoparticles must be functionalized in a way that guarantees efficient targeting without sacrificing the carrier’s overall structure.

The limitations of each component alone can be addressed by combining these materials. The combination of chitosan, PCL, PEG, spermine, and folic acid in a nanocarrier system may optimize the therapeutic advantages of gene delivery. For example, PCL can offer structural integrity and sustained release properties, whereas chitosan can supply the cationic charge required to condense DNA or RNA. Spermine can promote effective cellular uptake, whereas PEG can lengthen the circulation time, aiding in immune response evasion. By improving localization to cancer cells and possibly minimizing off-target effects, folic acid can guarantee targeted delivery. Previous studies have demonstrated the effectiveness of such multicomponent formulations. For example, Bakar et al. (2017) developed chitosan/PLGA nanoparticles for the delivery of DNA and reported improved transfection levels in cancer cells when these nanoparticles were combined with folic acid for targeted delivery. Similarly, research conducted by Jin et al. (2023) revealed that the integration of spermine in PCL nanoparticles significantly enhanced gene delivery efficiency by promoting endosomal escape. In conclusion, the synergistic qualities of chitosan, PCL, PEG, spermine, and folic acid nanoparticles suggest that their combined use could increase the effectiveness of gene delivery. This multifaceted approach may result in more effective therapeutic strategies for a variety of diseases by utilizing the benefits of each individual component while minimizing its drawbacks. This is especially true for cancer treatment, where targeted genetic material delivery is crucial.

Gene transfer is more sensitive than chemotherapy drug transfer since DNA is larger and more susceptible to damage during the process [42]. The immune system often recognizes and degrades nucleases, which can hinder delivery. Additionally, a successful gene delivery system must enable nanoparticles to release DNA without causing damage. Electrostatic interactions between cationic polymers and DNA neutralize the latter’s negative charge, reducing repulsion with the cell membrane and promoting stable DNA packaging into spherical structures [8]. These interactions increase gene delivery efficiency while protecting DNA from restriction enzymes and other harmful agents in the bloodstream. For restriction enzymes to digest DNA, they must bind to specific sites, but the coating of cationic polymers saturates the DNA surface, preventing enzyme access [52]. As shown in **Fig. 8**, micelles have significant potential in protecting DNA from plasma restriction enzymes.

The rapid release of drugs into plasma before they reach target tissues poses a significant challenge for drug delivery systems [53]. Liposomes, which are hydrophobic and cationic nanoparticles, are effective for *in vitro* gene transfer but have limited use in gene therapy because of the quick release of DNA into plasma [54]. This instability occurs because hydrophobic proteins and polyanions in plasma interact with the surface of the liposome, leading to premature DNA release.

Our studies indicate that surface modification of nanoparticles with PEG can prevent rapid DNA release under physiological conditions (pH = 7.4) (**Fig. 9**), which aligns with the findings of Abebe et al. [55]. After reaching target tissues, micelles enter cells via endosomes, but much of the DNA taken up is subsequently recognized and removed by lysosomes [55]. Therefore, releasing DNA from the endosome before it integrates with the lysosome is critical. Our findings revealed that DNA is released more rapidly from nanoparticles under acidic conditions, which may be due to the faster degradation of PEG in such environments. Furthermore, the proton sponge effect induced by cationic polymers can facilitate DNA release prior to lysosomal integration within the endosome.

Several approaches can be utilized to regulate the release of DNA from micelles, one of which involves adjusting the molecular weight of the hydrophobic components within nanoparticles (NPs) [6]. The incorporation of a hydrophobic polymer such as PCL into the NP structure enables this control. As shown in **Fig. 9**, attaching PCL to spermine regulated DNA release under neutral conditions but did not prevent release under acidic conditions (**Fig. 9**).

Magnetic nanoparticles can enhance gene and drug delivery systems by enabling imaging and targeted drug delivery through their accumulation in tumor tissues via an external magnetic field [56]. Additionally, metal nanoparticles can implement targeted hyperthermia by converting electromagnetic energy into heat [57]. Fe_3_O_4_ nanoparticles have strong magnetic properties, making them suitable for MRI and CT applications [45]. In this study, micelles were designed with magnetic nanoparticles incorporated in their surface layer, preserving the magnetic properties of Fe_3_O_4_. The conductive nature of Fe_3_O_4_ also facilitates heat transfer to the hydrophobic PCL layer of the micelles [58,59]. This heat transfer accelerates the degradation of the micelles, resulting in quicker DNA release into the tumor tissue [60].

The cytotoxic effects of certain nanocarriers are a significant barrier to effective gene delivery systems, highlighting the need for thorough evaluation of their toxicity. As depicted in **Fig. 10**, the chitosan-PCL/PEG-FA (CPPF) and spermine-PCL-chitosan/PEG-FA (SCCPF)/Fe_3_O_4_/DNA nanoparticles demonstrated negligible cytotoxicity, achieving over 80% viability in AGS cells. In contrast, free PEI was highly toxic to these cells. Other studies have reported similar observations. For example, Chang et al. reported that integrating PEI into PLGA nanoparticles markedly improved their cytotoxicity profile. The reduced cytotoxicity of the Spermine-PCL-Chitosan/PEG-FA/PEI-DNA nanoparticles may be attributed to their ability to shield the cationic charge of PEI, which helps prevent membrane damage caused by PEI [61].

As shown in **Fig. 12**, the transfection efficiency of PEI/DNA complex nanoparticles was significantly lower in serum-containing media than in SCPPF/Fe_3_O_4_/DNA. PEI/DNA nanoparticles have been widely used in various experiments because of their unique capabilities for nanoparticle delivery and easy integration with cell membranes. However, studies have shown that the efficiency of these nanoparticles is significantly reduced in serum-containing environments because of the presence of proteins in serum, which can lead to complex thermodynamics and changes in the structure of the nanoparticles, resulting in negative effects on their density and transfection efficiency [62]. In contrast, the SCPPF/Fe_3_O_4_/DNA nanoparticles performed better under similar conditions because of their unique composition and ability to tolerate various environmental conditions. These nanoparticles easily manage interactions with serum proteins and increase transfection efficiency [63].

To minimize nonspecific interactions between cationic nanoparticles and serum components, hydrophilic polymers such as PEG are often added to the nanoparticle surface [64]. PEGylation of polycationic/DNA nanoparticles helps prevent interactions with blood components and maintains their size [65]. Previous studies have indicated that using biodegradable cationic and amphipathic polymers together enhances the stability and gene delivery efficiency of nanoparticles [66].

Modifying nanoparticles with tumor-targeting receptors, such as folic acid and glucose receptors, represents a promising avenue for enhancing disease diagnosis and developing effective therapies [28]. Folic acid acts as a targeting ligand to improve the efficiency of drug delivery while minimizing adverse effects. However, the effectiveness of gene transfer is constrained when folic acid receptors on cancer cells become saturated. Increasing the concentration of nanoparticles beyond this point may lead to increased toxicity due to nonspecific uptake.

## Conclusion

This study aimed to develop micelles for efficient gene delivery to cancer cells while minimizing toxicity and side effects. We successfully synthesized SCPPF/Fe_3_O_4_/DNA micelles for targeted delivery to the AGS cell line. These micelles exhibited sustained DNA release in neutral environments and rapid release under acidic conditions, increasing drug availability in cancerous tissues while limiting release in normal tissues. This characteristic improves the efficacy of the nanoparticles on cancer cells and reduces their harmful effects on healthy cells. Furthermore, the SCPPF/Fe_3_O_4_/DNA nanoparticles demonstrated high stability in the presence of dextran sulfate at a neutral pH of 7.4. Our findings indicated that the micelles effectively shielded DNA from mechanical degradation and enzymatic digestion, facilitating the escape of internalized DNA from endosomes. As a result, the SCPPF/Fe_3_O_4_/DNA micelles achieved significantly greater transfection efficiency than did the PEI/DNA complexes. Overall, our results suggest that SCPPF/Fe_3_O_4_/DNA micelles represent a promising option for safe and effective cancer gene therapy, offering superior gene transfection efficiency, reduced cytotoxicity, and enhanced serum compatibility compared with those of the PEI/DNA formulation.

## Abbreviations

MTT: 3-(4,5-Dimethyl-2-thiazolyl)-2,5-diphenyl-2H-tetrazolium bromide
DMEM: Dulbecco’s modified Eagle’s medium
DLS: Dynamic light scattering
FBS: Fetal bovine serum
FA: Folic acid
^1^H-NMR: Hydrogen nuclear magnetic resonance spectroscopy
FTIR: Infrared spectroscopy
Fe_3_O_4_: Iron oxide nanoparticles
DMSO: Methyl sulfoxide
NPs: Nanoparticles
NCBI: National Cell Bank of Iran
OA: Oleic acid
PCPF: PCL-chitosan-PEG-FA
PLGA: poly(lactic-co-glycolic acid)
PCL: Polycaprolactone
PDI: Polydispersity index
PEG: Polyethylene glycol
PEI: Polyethylenimine
PLA: Polylactic acid
PVA: Polyvinyl alcohol
Que: Quercetin
RES: Reticuloendothelial system
SEM: Scanning electron microscopy
SP: Spermine
PCS: Spermine-chitosan-PCL
SPCPF: Spermine-PCL-chitosan-PEG-folic acid
TGA: Thermogravimetric analysis
TEM: Transmission electron microscopy
VSM: vibrating sample magnetometer

## Author contributions

MN and NB conducted the experiments, analysed the data, and wrote the original draft. HY and SM conceived and designed the research, administered and supervised the project, and reviewed and edited the manuscript. SB collaborated in the implementation of the molecular analysis and flow cytometry and reviewed the final version of the manuscript. All the authors read and approved the manuscript.

## Ethical statement

This article does not contain any studies with human participants or animals performed by any of the authors.

## Data availability

The authors declare that the data supporting the findings of this study are available within the paper. Should any raw data files be needed in another format they are available from the corresponding author upon reasonable request

## Consent for publication

Not applicable

## Participating declaration

Not applicable

